# Paleochronic reversion in *Psophocarpus*: Dynamics II, rhizogeny on floral anatomic fields

**DOI:** 10.1101/253955

**Authors:** Edward. G.F. Benya

## Abstract

Paleochronic reversion is confirmed in *Psophocarpus* as a basic floral ground state. That state can expand to include dynamics (T_(g,…,h)_) of axial expansion (AE) as a permutation (T_x_) phase beginning as phyllotactic floral phylloid (T_phyld_) and/or axial decompression (T_Axl_) manifest as linear elongation (T_Long_) and/or rotation (T_Rtn_) and/or latitudinal (T_Lat_) expansion. Organ regions present a continuum as a vector space 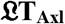 of floral axial transformation. A generative phase of meristem activity (T_(Rz, Sam, Infl)_) can follow.

Experiments with 49 phylloid and/or phyllome paleochronically reverted flowers presented varying degrees of phyllotactic permutation involving development of a pericladial stalk (PCL) and/or inter-bracts stem (IBS) and/or activated pedicel (PdcL) and/or gynophore (Gnf), Cupule-like (Gnf)/Cupl-Lk) elongation. A meristem generative function included rhizogeny as root site generation (RSG) at the calyx (Cl), PCL, bracts (Bt), IBS, PdcL and/or Gnf/Cupl-Lk regions manifest as eigenvector functions as RSG whose density of generation was associated with permutation of the ground state. A continuum of pedicel to calyx regions constitutes a subset 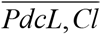 of a linear vector space 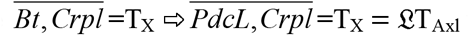 whose extension is defined within the space: 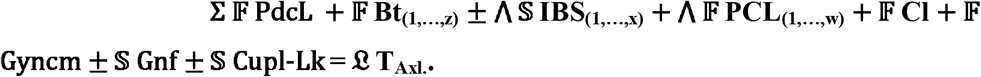.

The vector space 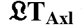 transforms to a master vector field 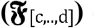 of generated Euclidian eigenvectors so that:

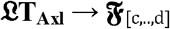

**Highlights:**

- Paleochronic reversion in plants presents paleo-botanic floral characteristics at a morphologic ground state.
- A permutative dynamic can ensue changing floral architecture.
- That dynamic can then transform to meristem generation presenting rhizogenous eigenvectors.

## Introduction

Angiosperm reproductive systems (Fig. 1) present five groups of organs in two anatomic zones; one pre-whorls zone of two region: pedicel and bracts, and an anatomic whorls zone of four concentric floral whorls regions: sepals, petals, stamens and ovary, all being usually well specified genetically and morphologically distinct (Stein, 1988; Kramer and Irish, 1999). Homeosis, as qualitative structural and/or positional variation, has been documented on numerous angiosperm species (Benya, 2000; Tookey and Battey, 2000; Kapoor, *et al.* 2002; Douglas and Riggs, 2005; Song and Clark, 2005). Such variation can affect organs or organ systems (Schwarz-Sommer, *et. al.* 1990; Pelaz, et. al. 2000), meristem identity (Ikeda, *et. al.* 2005), permutation (Coen and Meyerowitz, 1991; Parcy *et. al.* 1998; Benya, 2012) or body plan of an organism. Homeosis can also involve spatiotemporal patterns (Bäurle and Dean, 2006) affecting chronologies as homeotic conversion (Lenhard *et. al.* 2001; Jack, 2004) or reversion (Okamuro *et. al.* 1996; Honma and Goto, 2001; Parcy *et. al.* 2002).

**FIG 1.**
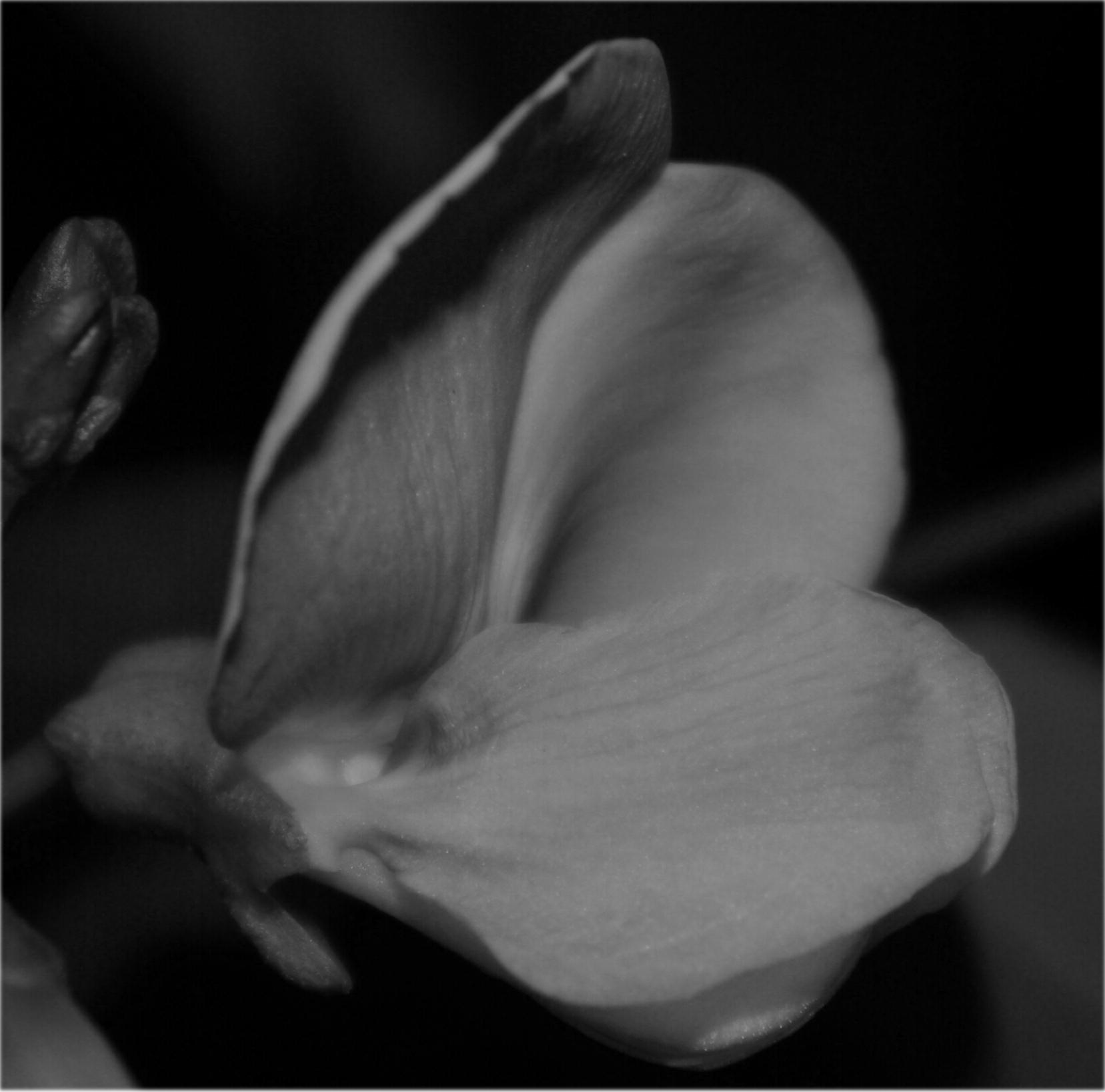
Flowers *Psophocarpus tetragonolobus*; normal (i.e. wild type), sexually reproductiave.

Homeosis in phyllome ground state floral specimens of *Arabidopsis* presents an “ideal basic organ” postulated by J.W. Goethe at the “foliar ground state” (Weigel and Meyerowitz, 1994). It can further involve development or expansion of nascent structures (e.g. gynophore) (Ditta *et. al.* 2004) and/or organ boundaries (Gendron *et al* 2012). A “phylloid ground state” (Schwarz-Sommer, *et. al.* 1990) of floral specimens of the species *(Psophocarpus tetragonolobus* (L.) DC), variable in form (Benya and Windisch, 2007, Fig. 2), could then present further development of nascent structures (e.g. pericladial stalk) (Fig. 3) and/or organ boundaries. This gave rise to morphological spacing of the floral whorls zone from the pre-whorl pedicel-bracts zone thus “distancing” (McLean and Ivimey-Cook, 1961) both anatomic zones from each other through phyllotactic floral axial permutation (Benya and Windisch, 2007). Goal of this research was to test paleochronically reverted flowers (i.e. experimental specimens) at a phylloid and/or phyllome ground state of the species *P. tetragonolobus* (L.) DC (Fabaceae) for any meristematic generative capacity (e.g. rhizogenous site genesis - RSG, shoot apical meristem - SAM).

**FIG 2.**
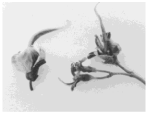
Reverted flower and inflorescence presenting beginning decompression permutation (webbing of carpel).

**Fig. 3.**
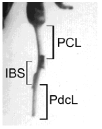
Bract dislocation demonstrating a pericladial stalk (PCL) of about 15 mm, an “inter-bractial stem” (IBS) of about 12 mm on a pedicel (PdcL) (partial) of more than 12 mm.

## Materials and methods

Research examined 49 paleochronically reverted flowers (i.e. experimental specimens, Table 1) at a phylloid and/or phyllome ground state (Benya, 2016; Fig. 1 therein) originating from field-grown homeotic segregants (i.e. homozygous, recessive recombinants) (Benya, 1995) of the species *P. tetragonolobus* (L.) DC (Fabaceae). Specimens presenting bracts and whorls anatomic regions juxtaposed or those presenting phyllotactic alteration (e.g. “permutatively distanced” [Benya, 2012]) were tested for any meristematic generative capacity (e.g. root site genesis - RSG and/or shoot apical meristem - SAM genesis). SAM’s were identified by their shoot structure. Rhizogeny was also identified by structure; white growth structures being assumed to be rhizogenous according to Aida *et al.* 2004. Meristematic sites were identified qualitatively by location within any of six morphologic regions; pedicel (PdcL), bracts (Bt), pericladial stalk (PCL), calyces (Cl), gynophore (Gnf) and/or cupule-like (Cupl-Lk) structure. Within the whorls zone, at the conjunction of the androecium and the gynoecium, a gynophore and/or cupule-like structure could arise (henceforth “gynophore/cupule-like” or Gnf/Cupl-Lk), presented as such because the homology of both structure is no yet confirmed.

**Table 1.**
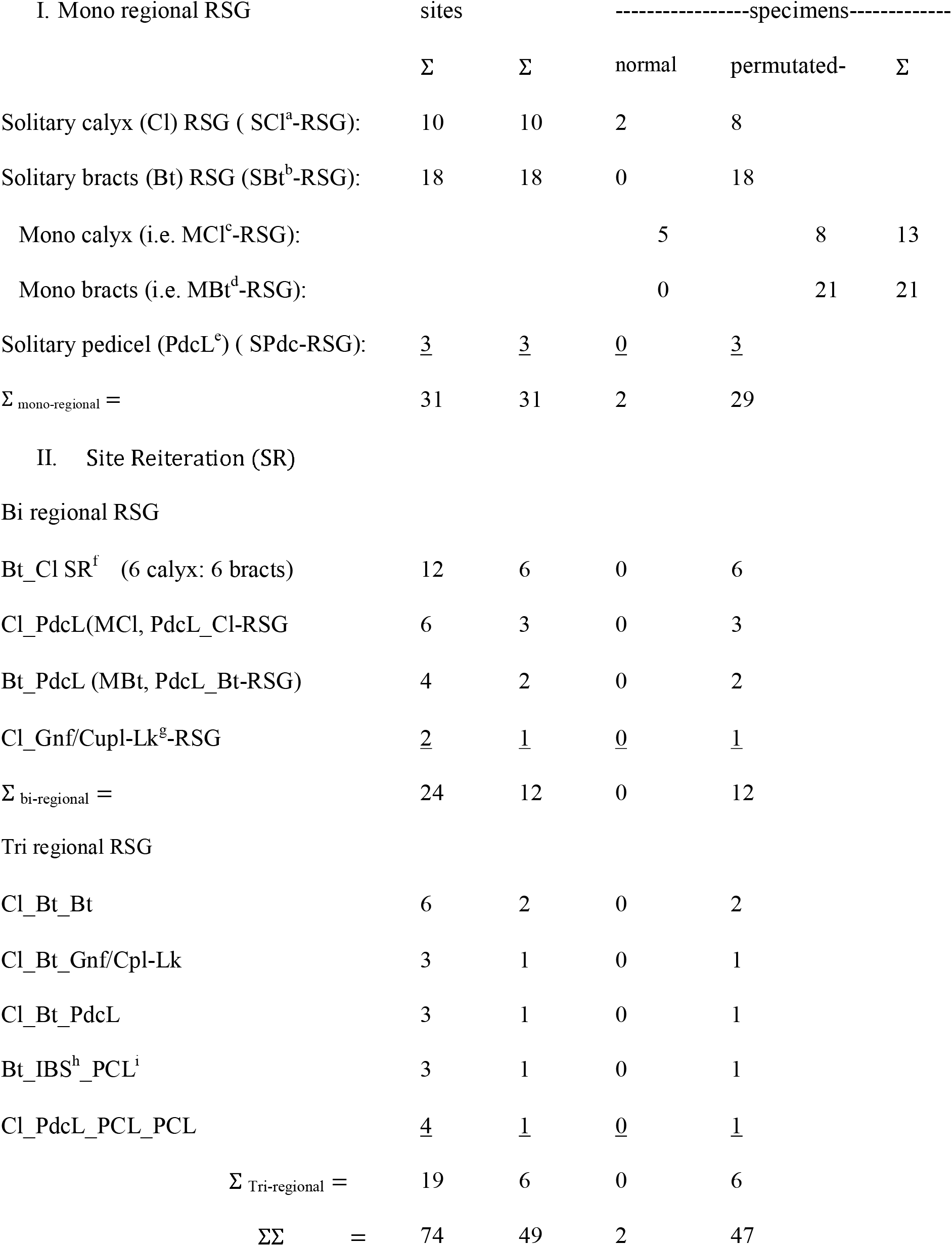
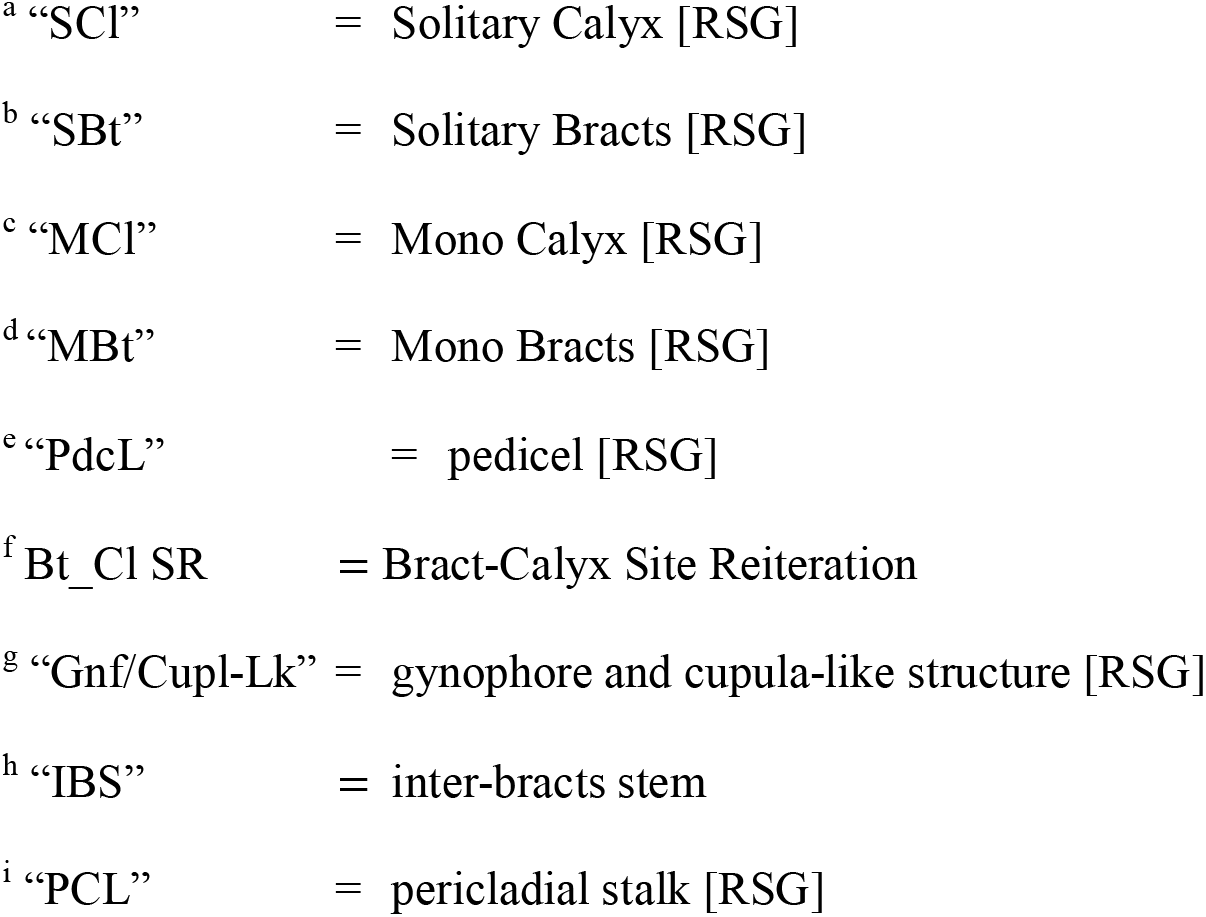
The 49 specimens presented a multiplicative regional distribution of:

| I. Mono regional RSG | sites |  | -----specimens----- |  |  |
| --- | --- | --- | --- | --- | --- |
| | $\Sigma$ | $\Sigma$ | normal | permuted- | $\Sigma$ |
| Solitary calyx (Cl) RSG ( $SCI^a$-RSG): | 10 | 10 | 2 | 8 | |
| Solitary bracts (Bt) RSG ( $SBt^b$-RSG): | 18 | 18 | 0 | 18 | |
| Mono calyx (i.e. $MCl^c$-RSG): | | | 5 | 8 | 13 |
| Mono bracts (i.e. $MBt^d$-RSG): | | | 0 | 21 | 21 |
| Solitary pedicel (PdcL<sup>e</sup>) ( $SPdc$-RSG): | <u>3</u> | <u>3</u> | <u>0</u> | <u>3</u> | |
| $\Sigma_{\text{mono-regional}} =$ | 31 | 31 | 2 | 29 | |
| II. Site Reiteration (SR) |  |  |  |  |  |
| Bi regional RSG |  |  |  |  |  |
| Bt_Cl SR<sup>f</sup> (6 calyx: 6 bracts) | 12 | 6 | 0 | 6 |  |
| Cl_PdcL($MCl$, $PdcL\_Cl$-RSG | 6 | 3 | 0 | 3 | |
| Bt_PdcL ($MBt$, $PdcL\_Bt$-RSG) | 4 | 2 | 0 | 2 | |
| Cl_Gnf/Cupl-Lk<sup>g</sup>-RSG | <u>2</u> | <u>1</u> | <u>0</u> | <u>1</u> |  |
| $\Sigma_{\text{bi-regional}} =$ | 24 | 12 | 0 | 12 | |
| Tri regional RSG |  |  |  |  |  |
| Cl_Bt_Bt | 6 | 2 | 0 | 2 |  |
| Cl_Bt_Gnf/Cpl-Lk | 3 | 1 | 0 | 1 |  |
| Cl_Bt_PdcL | 3 | 1 | 0 | 1 |  |
| Bt_IBS<sup>h</sup>_PCL<sup>i</sup> | 3 | 1 | 0 | 1 |  |
| Cl_PdcL_PCL_PCL | <u>4</u> | <u>1</u> | <u>0</u> | <u>1</u> |  |
| $\Sigma_{\text{Tri-regional}} =$ | 19 | 6 | 0 | 6 | |
| $\Sigma\Sigma =$ | 74 | 49 | 2 | 47 | |

- <sup>a</sup> “SCI” = Solitary Calyx [RSG] - <sup>b</sup> “SBt” = Solitary Bracts [RSG] - <sup>c</sup> “MCI” = Mono Calyx [RSG] - <sup>d</sup> “MBt” = Mono Bracts [RSG] - <sup>e</sup> “PdcL” = pedicel [RSG] - <sup>f</sup> Bt\_Cl SR = Bract-Calyx Site Reiteration - <sup>g</sup> “Gnf/Cupl-Lk” = gynophore and cupula-like structure [RSG] - <sup>h</sup> “IBS” = inter-bracts stem - <sup>i</sup> “PCL” = pericladial stalk [RSG]

Quantitative count of meristem site numbers per specimen then furnished a measure of their intensity and distribution. Statistical analysis (SPSS, 2013) served to test for significant variation in meristem distribution that might be associated with phyllotaxis and/or phyllotactic alteration specifically between the whorls anatomic zone, whose lowest boundary is that of the sepals (i.e. calyx) region, and the non-whorls floral zone defined by the pedicel and bracts regions.

Specimens distributed over 21 treatments containing one to four reps per treatment primarily oriented to determining any age and/or timing variables influencing meristematic activity. Treatments were conducted in individual glass test-tubes and/or plastic cups in plain or deionized water within a simple-structured laboratory. Humidity and temperatures within the laboratory followed closely those of external ambiental conditions during the winter and spring seasons in the semi-arid, tropical equatorial climate of Teresina, Piaui, Brazil (05^o^05’S; 42^o^49’W, alt. 64 m) from early August to late November (Gadelha de Lima, 1987). The laboratory had no internal climate control.

Seven morphologic categories served to classify results by physical location. The first and second categories involved bracts (Bt) “juxtaposed” to the calyx (Cl) (i.e. normal, non-permutated), thus recognizing meristem genesis at bracts and/or calyx loci. The third and fourth categories involved phyllotactically altered specimens whose floral axes were permutated (Benya, 2016). These presented bracts that were physically “distanced” from the calyces by a pericladial stalk (PCL) and, at times, occasional bract dislocation due to development of an inter-bracts stem (IBS, Fig. 3) (Benya and Windisch, 2007) or, at times, on the pedicel (PdcL) below the last (lowest) bract (Douglas and Riggs, 2005). Floral axial permutation permitted recognition of anatomic zones, morphologic regions and loci distinguishing organ regions from normal anatomic organ loci of non-permutated specimens. Loci of the bracts anatomically defined the bracts region which extended from zero (i.e. bracts parallel or normal) to 12 mm depending on development of any IBS. Specimens showing only a single bract (having lost one) received a measurement from that one bract to the calyx.

Any change in phyllotaxis for a specimen usually occurred during the field stage. Thus each specimen entered its respective treatment with a test phyllotaxis usually well established. This presented a significant degree of stability (Benya, 2016). Quantitative distribution of the 49 specimens was: two having no change and 47 showing some form of phyllotactic change, but 43 presenting pericladial stalks and six of the 49 being “normal” (juxtaposed calyx-bracts regions).

The fifth category of potential meristem genesis, the pericladial stalk (1 to 38 mm), was treated as a distinct morphologic region in itself. Because the lengths of the stalks were basically stable but more than one specimen could present the same length (e.g. three specimens presenting stalk lengths of 11.0 mm) (Supplementary data, Fig 1), the varying numbers of specimens, presenting the same stalk lengths formed natural clusters of millimetric intervals. The sixth category of possible meristem genesis could be at any pedicel segment below the lowest bract (i.e. the pedicel or part thereof). Thus permutated pedicel distance (meristematically active pedicel length), distinct from non-permutated distance, was assumed to be the measure from any RSG locus on the pedicel to the lowest bract.

The seventh category could be at the gynophore and/or cupule-like structure. The term “and/or” used herein is because RSG arose at the node of the convergence of the gynophore with the cupule-like structure. Qualitatively confirmed permutated regions (e.g. gynophores, cupule-like, activated pedicel) but whose exact measurement could not be confirmed, received a measurement of one (“1.0”) mm for purposes of statistical analysis.

Analysis then proceeded to measure any vectorial biophysical functions. Because of complexities involving qualitative and quantitative aspects of the physiological basis of rhizogeny (Bao, *et al.* 2007) and references cited therein; (Müller and Sheen, 2008), this report addresses only the biophysical aspects of meristem genesis.

## Results

No SAM generation occurred. Rhizogeny as “root site generation” (RSG) occurred at any of six morphologic regions (i.e. pedicel, bracts, pericladial stalk, calyx, gynophore/cupula-like structure) as 74 active sites, primary and confirmatory, were distributed among the 49 specimens (Table 1) (1.5 sites per specimen).

Further RSG might arise, but mainly for reasons of temporal/spatial dynamics, these analyses focus on initiatory sites with attention to a secondary PCL and five bracts loci on three specimens (one at each bract and one IBS; technically three of which are secondary sites but are treated here as primary for purposes of exactitude). This helped distinguish a more general cadastral generative dynamic (Weigel and Meyerowitz, 1994) from the specific vectorial cadastral (Fiehn *et al.* 2000) dynamic.

At calyx-bracts juxtaposition, RSG was recognized as originating from the calyx if the root emerged from the lower-fusion (connation) of the calyx or above that fusion (five specimens). It was recognized as originating from the bracts if the root emerges from the conjunction of the pedicel with the calyx (none).

Five of six “normal” specimens (i.e. bracts-calyx regions juxtaposed) presented RSG at their calyces. Three of the six had RSG only at the calyces (solitary-calyx; SCl RSG) two had RSG at calyx (mono-calyx; MCl RSG) plus pedicel (PdcL). One had RSG only on the pedicel solitary PdcL (i.e. SPdcL RSG). Two of the 47 phyllotactically permutated specimens presented RSG on their pericladial stalks (Fig. 4) plus other sites, complementary-site PCL (CoS-PCL) RSG.

**Fig 4.**
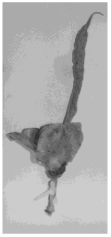
Primacy of RSG on pericladial stalk.

Thirty one specimens presented single (i.e. mono-regional) RSG (i.e. 31 MR-RSG). Ten specimens presented solitary-regional organ site RSG only at their calyces (10 SCl-RSG). Eight of these were permutated at non-calyx-bracts regions and two were completely without permutation: normal. Three other specimens with permutated calyx-bracts regions, had calyx plus non-bracts regional RSG; mono site calyx (MCl) RSG (Table 1).

Twenty one (21) specimens, all permutated showed mono-regional organ site RSG only at the bracts. One of these included distinct bract, IBS and PCL RSG loci. Two others had bract and pedicel RSG yielding a total of three “mono bract” (3 MBt-RSG) specimens leaving 18 “solitary bracts” (SBt) RSG specimens. Three specimens presented mono-regional RSG only on the pedicel (solitary pedicel RSG or three SPdcL-RSG) (Table 1).

Eighteen (18) specimens, all permutated, showed reiterative RSG of two to four sites at two to three (i.e. “bi” or “tri”) anatomic regions per specimen. Ten of these presented singular bracts-calyx site reiteration (Bt-Cl SR) RSG; six at both bracts and calyx; bi-regional Bt_Cl reiteration (Bi Cl_Bt SR-RSG). Six others had diverse organs, non-organs bi-regional RSG; three at calyx and pedicel, two at bracts and pedicels, and one at its calyx and gynophore/cupule-like regions; (“bi-regional SR” or 6 BiSR-RSG) (Fig. 5) (Table 1).

**Fig 5.**
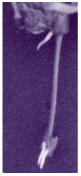
RSG at both bracts and calyx organs regions “biregional distribution” of 3 rhizogenous sites or “OSR-RSG” (i.e. “organ site reiteration”) about 15 cm between calyx and bracts.

Six specimens presented tri-regional RSG: one at its calyx, bracts and gynophores/cupule-like regions; one at its PCL, at a bract, and in its IBS region; one at its calyx, bracts and pedicel regions; and two at their calyces, but then one at each bract, the bracts regions having been permutated by an IBS. The sixth specimen had tri-regional RSG at four loci: one on the pedicel, one at its calyx, and two at the pericladial stalk, distinct in time and loci for purposes of timing (Table 1).

The 49 specimens presented a multiplicative regional distribution of the 74 RSG sites (Table 1). RSG at the three organs regions (Bt, Cl, Gnf/Cupl-Lk) totaled 60 sites: 25 at calyces and 33 at the bracts and two at the gynophore/cupule-like regions. However, 58 of these 60 were significantly concentrated (F_3,45_ =153.781, p<0.000) at the calyx-bracts organs regions (Table 2). RSG at the bracts and calyx sites was significantly above RSG at all other anatomic sites (F_3,45_ =15.577, p<0.000). This significance (58 of 74 sites) defined these two regions, anatomically and morphologically as spatio-temporal centers of meristematic activity. Primary site generation between both organ regions ran significantly low at the calyces and high (r= – 0.360, p= 0.011) at the bracts (ANOVA, F_1, 47_ =6.980, r^2^ = 0.129). RSG intensity between the group of 10 SCl-RSG specimens and the 21 MBt-RSG specimens (Table 1) presented further negative linear correlation (r = – 0.423) and dynamics (r^2^ = 0.179, F_1,44_ = 9.565, p = 0.003), significantly different intensities of RSG generative dynamics for each organs region. This permitted a basic vectorial (i.e. eigenvector) approach (Fiehn et al. 2000) to analyses wherein vectors were collectively recognized as well as any scalar functions (λ) and intensities indicative of one or more eigenvectors (*E^n^*[*v*]_(1,…,n)_).

**Table 2:** RSG at the two principal organs regions, 23 at calyces (Cl) and 35 at the bracts (Bt), totaled 58 sites in a distributive site ratio on 46 of the 49 specimens as:

| RSG | Mono regional |  | <u>Bi regional:</u> |  | <u>Tri regional:</u> |  | <u>Σ</u> |  |
| --- | --- | --- | --- | --- | --- | --- | --- | --- |
| Organs: | Cl | Bt | Cl | Bt | Cl | Bt | Cl | Bt |
| Sites (Cl distribution): | 15 |  | 8 |  | 2 |  | 25 |  |
| Sites (Bt distribution): |  | 21 |  | 8 |  | 4 |  | 33 |
| Σ Sites: | 15 | 21 | 16 |  | 6 |  | 58 |  |
| Σ Specimens: | 15 | 21 | 8 |  | 2 |  | 46 |  |

Quantitatively the PCL, IBS dislocation and PdcL define a physical continuum of distance(s) or spacing between calyces, bracts and pedicel but the pericladial stalk dominates, present on 42 of the 47 permutated specimens. Thus it is used as the principle measure of floral axial expansion (AE). Physical axial continuum (AC) of the gynophore/cupule-like regions with the calyx, pericladial stalk and regions of the pre-whorls zone, as in previous research (Benya, 2016), cannot be verified by this data.

However an axial generative continuum (AGC) of RSG runs from the PdcL to the Cupl-Lk structure inclusive.

RSG at those organs and regions was analyzed statistically in terms of significant dependent and independent relations (ANOVA), correlations and dynamics (multiple regressions) with axial lengths (e.g. PCL). Invariability or minimal variability of such analyses up to and including proportional repetition or diminution of results from different samples (e.g. organs sites, bi-regional, specimens’ meristem genesis, etc.) then revealed scalar(s) functions (λ) and intensities indicative of one or more eigenvectors (*v*).

Bract dislocation due to IBS development extended the linear intra-floral transformative function presented by floral axial permutation forming a biophysical field (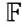 PCL) and subfield (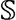 IBS). In a sample of six specimens presenting bract dislocation, all six showed RSG initially at the lowest bract (i.e. the one furthest from the calyx), or on the pedicel just below that bract indicating a field continuum (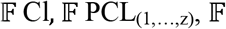 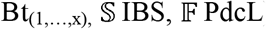) that depended on PCL and /or IBS and/or PdcL permutation.

Differences in the distribution and intensity of RSG were significantly related to time-age variables (in days). A significant negative linear correlation arose between length of total axial permutation and timing of initial RSG at any anatomic region dated from the beginning of each treatment (r= – 0.335, p=0.018, df = 47). A significant positive cubic regression arose between “rooting time since reversion” (i.e. “reversion age” of a specimen; time from initiation of reversion on a recombinant to the date when a reverted floral specimen from that recombinant was collected for study, plus the time of initial RSG on that specimen) and, in sequence, the “timing of initial RSG” at any anatomic region on that specimen (cubic r^2^ = 0.199, F_3, 45_ = 3.738, p = 0.018). This indicated a “capacitation time” necessary for the RSG function to arise on a specimen counting from initiation of reversion; about 56 days.

Significant floral linear (651 mm) decompression (r^2^ = 0.099, F_1, 47_ = 5.187, r= 0.315 p = 0.027), confirmed the overall advantage in RSG (74 sites) imparted by floral axial permutations. That permutation verified a linear progression and amplification of cadastral RSG eigenvector function from the calyx regions to the bracts regions.

Intensity of RSG on specimens presenting bi-regional Bt_Cl reiterative RSG (22 sites on 10 specimens) was significantly correlated with permutation of the floral axis (651 mm) over time (r = 0.387, p = 0.008); reflected in the calyx-bracts bi-regional plastochron. Transition (T) of 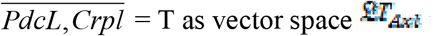 as vector space 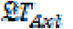 to vector field 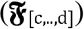 presents a dynamic of extension and transformation of function from; axial expansion permutation to a generative function (e.g. RSG) that is manifest as Euclidian eigenvectors.

The most obvious indicators of eigenvectors emerged from data for the 10 SCl-RSG specimens and the 21 SBt-RSG specimens. The most striking eigenvector indicators however came from a comparison involving combined RSG values for the six specimens (8 sites) presenting juxtaposed calyx-bracts sites with the 10 permutated specimens presenting bract-calyx SR-RSG (Bt_Cl SR-RSG) whose sum of 10 calyx sites more than replicated to 12 bracts sites (Σ_sits_ = 22). Within that cluster of 10 specimens, seven presented calyx RSG whose initiation time preceded (was less than) the bract initiation time on those specimens. One presented equal calyx and bract RSG initiation time. Two specimens presented RSG at four distinct bract sites; at two bract loci each distanced by an IBS, two at each specimen. These preceded calyx RSG (Table 3).

**Table 3:**
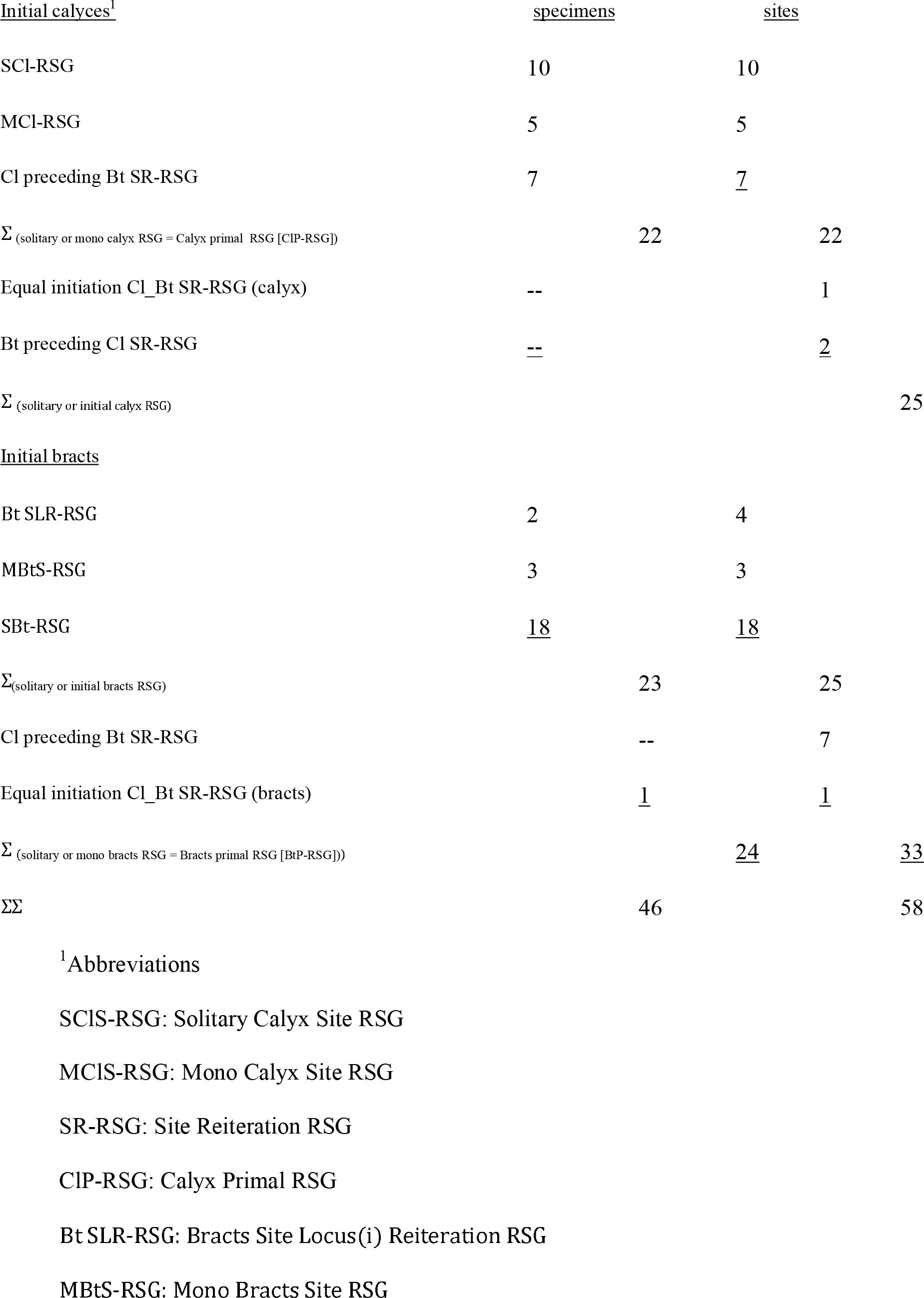
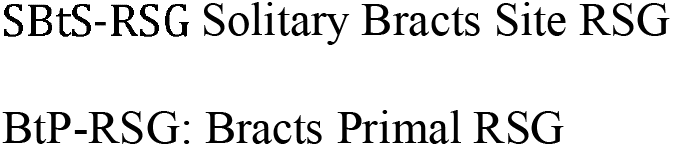
The sequence of calyx and bracts organ site generation.

| <u>Initial calyces</u><sup>1</sup> | <u>specimens</u> | <u>sites</u> |  |
| --- | --- | --- | --- |
| SCI-RSG | 10 | 10 |  |
| MCI-RSG | 5 | 5 |  |
| CI preceding Bt SR-RSG | 7 | <u>7</u> |  |
| $\Sigma$ (solitary or mono calyx RSG = Calyx primal RSG [CIP-RSG]) | 22 | 22 | |
| Equal initiation CI_Bt SR-RSG (calyx) | -- | 1 |  |
| Bt preceding CI SR-RSG | -- | <u>2</u> |  |
| $\Sigma$ (solitary or initial calyx RSG) | | | 25 |
| <u>Initial bracts</u> |  |  |  |
| Bt SLR-RSG | 2 | 4 |  |
| MBtS-RSG | 3 | 3 |  |
| SBt-RSG | <u>18</u> | <u>18</u> |  |
| $\Sigma$ (solitary or initial bracts RSG) | 23 | 25 | |
| CI preceding Bt SR-RSG | -- | 7 |  |
| Equal initiation CI_Bt SR-RSG (bracts) | <u>1</u> | <u>1</u> |  |
| $\Sigma$ (solitary or mono bracts RSG = Bracts primal RSG [BtP-RSG]) | | <u>24</u> | <u>33</u> |
| $\Sigma\Sigma$ | 46 | | 58 |

<sup>1</sup>Abbreviations
SCIS-RSG: Solitary Calyx Site RSG
MCIS-RSG: Mono Calyx Site RSG
SR-RSG: Site Reiteration RSG
CIP-RSG: Calyx Primal RSG
Bt SLR-RSG: Bracts Site Locus(i) Reiteration RSG
MBtS-RSG: Mono Bracts Site RSG
SBtS-RSG Solitary Bracts Site RSG
BtP-RSG: Bracts Primal RSG

Mathematically distribution of the 58 Bt_Cl sites on 46 specimens is confirmed by a variable but predictable, significant linear (r^2^ = 0.129, F_1,47_ = 6.980, p = 0.011) transfer (r = – 0.360) and amplification (quadratic r^2^ = 0.323, F_2, 46_ = 10.960, p < 0.000) of eigenvector function. This dynamic began at the calyx region (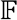 Cl), amplified to include the pedicel (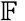 PdcL) regions then was confined to bracts and/or pedicel over the first 4.0 mm of PCL permutation. It amplified again to include Cl-RSG over the continuing floral permutation, an overall cubic regression (r^2^ = 0.161, F_3,45_ = 2.884, p < 0.046). This dynamic was selective, preferential in distribution and manifestation, but neither exclusive nor random, running at a general scalar of (λ = 1.3); summarized as:

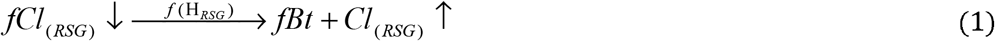

The two completely normal specimens (no permutation) presented solitary RSG only at their calyx regions. As overall permutation began (i.e. Gnf/Cupl and/or pedicel regions), solitary RSG arose at the calyx of one specimen and pedicel of another. Solitary pedicel permutation also supported bi-regional RSG on two specimens, both (Cl_PdcL). Bracts-calyx regions remaining juxtaposed on all four specimens.

As PCL permutation began RSG regressed to the solitary condition at the bracts (four specimens). Bi-regional Bt_Cl-RSG first arose on a specimen with 4.0 mm of PCL, regressing to mono Cl-RSG on one specimen and Bt-RSG on another, both with 5.0 mm of PCL. RSG amplified again to bi-regional Bt_Cl on a specimen having 1.0 mm of PCL and 10.0 mm of IBS, regressing to mono Cl or Bt RSG on 10 specimens over the next 5.5 mm of PCL plus IBS permutation (Supplementary material, Fig 1 “map”).

As PCL and IBS regional permutation continued, Cl_Bt reiteration then reached a maximum intensity (four specimens) in a group of 10 specimens presenting 11 to 15 mm of PCL, but up to 17 mm when including 2.0 mm of IBS for one specimen. A total of 17 of the 74 RSG (22.97%) arose within that four mm permutation span; 1 PdcL 7 Bt, 7 Cl, and 2 Gnf/Cupl. Total organ-site RSG (60 of 74 sites) as well as bracts-calyx site reiteration, (22 sites on 10 specimens) followed normal distributions as lengths of PCL increased.

The six normal specimens, presenting eight RSG regions, constituted a cluster. These served as a reference both qualitative and quantitative to and for succeeding zones, areas, spaces, loci and regions where RSG presence, in Euclidean spaces 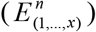 and eigenvector generation intensity could be manifest (e.g. 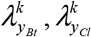). In simplest terms the pericladial stalk, bracts site (including IBS) and pedicel, constitute biophysical anatomic fields 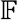 PCL, 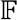 Bt_(1,…x)_ and 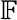 PdcL, a continuous morphologic physical avenue (i.e. vector space) of transport for the putative morphogen(s) and/or signal(s) (Roberts and Friml, 2009; Lohmann, *et. al.* 2010) responsible for RSG. These fields were essential for linear transport and/or transfer of the RSG function (*H*_(RSG)_) from one coordinate system (e.g. calyx) to another (e.g. bracts). Qualitatively the “path calyx” or pericladial stalk defined spatio-distancing between the two organs.

At permutation RSG progressed at a rate (*^r^RSG*_(0.0,…,*m*)_) that generally increased in intensity (Δ*k*_(>0.0,…,*m*)_) over the first 18 mm of PCL development but overall was exponentially variable where:

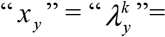 the number of specimens within pericladial stalk interval of ≥1.0 mm at exponential rates of: *k*_(0.0,…,*m*)_+Δ*k*_(>0.0,…,*m*)_.

This was reflected in significant quadratic regression (r^2^ = 0.123, F_2,46_ = 3.214, p = 0.049)

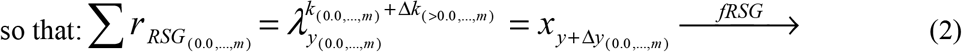

The concentration of Bt_Cl OSR RSG, essential to the generally increasing rate of RSG intensity, already mentioned, reached its maximum at 18.0 mm of stalk length. This reflected a definite tendency toward supplementarity of rhizogenous canalization. Of note was one specimen with a 30.0 mm PCL. It presented four distinct rhizogenous loci in a tri-regional arithmetic field distribution of 1:2:1; at Cl: PCL: and PdcL respectively (Table 1), thus presenting putative non-vetorial cadastral (Weigel and Meyerowitz, 1994) (i.e. PCL) as well as vectorial cadastral (Fiehn *et. al.* 2000) (i.e. Cl, PdcL) dynamics of RSG.

This generative sequence further supports the conjecture (Benya and Windisch, 2007) that morphologically distinguishes a true pericladium (McLean and Ivimey-Cook, 1961) from a pericladial stalk, the stalk quite possibly being a morphological extension of the calyx. The first of the PCL-RSG sites arose on a specimen where it preceded RSG at the calyx and the pedicel. The second was the last of the four initiatory RSG’s for that specimen.

Thus the 18 specimens presenting reiteration in a bi-regional to tri-regional (organs to organs and/or organs to stalk distribution) 12:6 ratio (Table 1), permitted sequential timing analysis (in days) for RSG intervals in an n-dimensional coordinate system of temporal [floral axial permutation] vector function with that of spatial RSG (eigen) vector field function. This complemented the spatial analysis. Considering reiteration only at organs regions (i.e. “organ site reiteration”: 11 OSR-RSG), initiation was manifest eight times at calyces, twice at bracts, once at equality (same day at bracts and calyces), and never initially at the gynophores and/or cupule-like structure, for a 8:2:1:0 ratio, a significant deviation 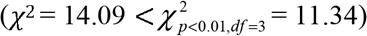 from random expectation.

Overall mean spacing between calyx-bracts sites (i.e. PCL permutation) was 11.33 mm. However it reached 13.21 mm on 42 permutated specimens presenting PCL permutation (range 1.00 to 38.0 mm).

The 10 calyx sites on 10 specimens showing SCl-RSG (eight of which were permutated with a total of 76 mm of pericladial stalk) presented mean distancing 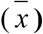 between Cl_Bt regions of 7.60 mm (range 0.00 to 21.00). Mean distance between organs regions increased to 11.81 mm (248/21), (range 1.0 to 32.00) for the 21 SBt-RSG specimens. As overall PCL permutation began and progressed, a significant linear shift (**r** = -0.680) occurred (r^2^ = 0.462, F_1, 47_ = 40.337, p < 0.000), in overall RSG focus, shifting principally form Cl regions to Bt regions (*f H*_RSG_ → Bt RSG). PCL development had no significant influence on RSG for these 10 SCl specimens. Although PCL development showed positive linear correlation with RSG at Cl_Bt organs sites (r = 0.301, p = 0.035, df = 47), timing of organs’ sites RSG was negatively correlated, although not significantly (r = – 0.099, p = 0.644, df = 22), with PCL development. This verified that, at permutation (especially as it began, of short PCL lengths), the earlier and more intensive rate of RSG occurred at the bracts.

## Discussion

A phylloid ground state and/or various degrees of phyllome organ formation (Weigel and Meyerowitz, 1994; Pelaz, *et. al.* 2000) characterized all 49 experimental specimens (Fig. 2, 3). Floral meristem cancellation (Benya and Windisch, 2007) was anticipatory and essential to that state. After cancellation the gynophores and/or cupule-like, pericladial stalk, calyx and bracts organs regions plus the pedicel usually entered an active permutation decompression phase (Benya, 2016) where elongation of structures changed floral phyllotaxis (Benya and Windisch, 2007). Distancing of anatomic regions frequently gave more distinction to organ positions. Loci at these regions could easily change position while organ forms remained quite constant (Fig. 3). However organ regional definitions could become complicated. Measuring permutation distance at the pedicel was complicate by the fact that normal, non-reverted pedicel length ranged from 5.0 to 13.0 mm at 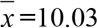 That length could increase to 20.0 mm in reversion.

The significant positive cubic regression between “rooting time since reversion” and the “timing of initial RSG” at any anatomic region on that specimen (cubic r^2^ = 0.199, F_3,45_ = 3.738, p = 0.018) verified a “capacitation” time (Benya, 1999) necessary for the RSG function to arise on a specimen counting from initiation of reversion; about 56 days. This included an average of about 46.5 days of permutation time, plus about 9.5 days for RSG. The dynamic of floral axial permutation as axial expansion (AE) was distinct from the dynamic of RSG. In this and previous studies (Benya, 2016), they never occurred concurrently nor showed any overlap. The two phases appear to be not only distinct but mutually exclusive.

Established morphologic differences between bracts and calyces emphasize anatomic qualities that define and distinguish them both (Stern, 1988). This research supports a premise of competitive interaction between the permutative and generative phases. Significant quadratic and cubic regressions further support a premise of competition between calyx and bracts organs at the RSG phase.

The organ regions; calyces, bracts (plus IBS therein) and gynophores/cupule-like structure, as vector fields 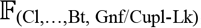, contain distinct euclidean eigenvectors (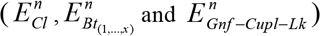 presenting temporal and physical sets having highly predictable capacities to generate and support RSG. For purposes of simplicity, anatomic regions are treated as two-dimensional euclidean spaces (e.g. *E*^2^) rather than as three-dimensional (*E*^3^) spaces, with no loss in mathematical accuracy.

The pericladial stalk serves as a qualitative anatomic structure whose varying morphologic lengths between each specimen, presented a biophysical field (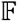 PCL) to distance the 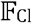 and 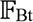 and the euclidean spaces therein from each other but not necessarily separate them. Supplementary RSG (as OSR or organ or non-organ RSG) is possible. The spatial sequence of RSG, beginning at calyces and running to bracts, although variable, was significant. This indicated the presence of a qualitatively distinct plastochron (a definite timing interval between initiation of RSG at one and then at the other organs region) (Lohman et. al. 2010). The anatomic regions are a succession of fields (i.e. 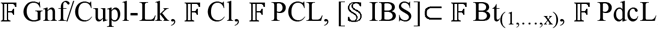) of the permutated floral axis (T_Axl_). These begin at the gynophores-cupule-like region. They then diminish or terminate spatially at the androecium and/or corolla. They arise again at calyx regions and extend to the pedicel inclusive, thus presenting two continua that are distinct spaces 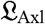.

Besides distancing bracts and calyces from each other, development of a pericladial stalk also distanced the entirety of the floral whorls anatomic zone from the pre-whorls (pedicel-bracts) zone. The generative phase at these areas and their respective regions was then characterized by a definite canalization (Okamuro *et. al.* 1993; Debat and David, 2001; Casci, 2005; Parkinson *et.al.* 2007) that was meristematically rhizogenous. Distribution of initial RSG, both between and within morphologic regions, presented variation significantly associated with phyllotactic permutation as axial expansion (AE).

Juxtaposition of Cl_Bt regions is the phyllotactic norm for these floral organs of this and most angiosperm species. Within the constraints of these investigations (i.e. considering the continuity of the whorls zone as one component, and the non-whorls pedicel-bracts zone as another), the calyx and/or pedicel were the primary regions of RSG at/or prior to any permutative distancing. The bracts then became the primary region of that activity after PCL distancing began and then the main focus of rhizogeny as permutation continued, but within predictable intensities and limits.

The pericladial stalk shows morphologic continuity with the calyx (Benya and Windisch, 2007) thus distinguishing it from a true pericladium (McLean and Ivimey-Cook, 1961) and raising questions of anatomic homogeneity. Highly significant statistical results support the hypothesis that the pericladial stalk is an anatomic area (a morphological extension) of the calyx organs region (Benya and Windisch, 2007). RSG on pericladial stalks while temporally quite predictable, was spatially not predictable. A putative, non-vectorial, cadastral generative function (i.e. mere specification of rhizogenous meristem identity) (Weigel and Meyerowitz, 1994) for RSG (or perhaps “leakage”), seems to be operative on the stalk.

The organs at both Cl and Bt regions maintain their specified identities at definite positions of their respective loci. However anatomic organ regional area expansion, by means of PCL and/or IBS regional development can augment organ regional dimensions and even change locus orientation and fields.

RSG followed significant timing and regional manifestation significantly concentrated at the three organs regions and especially at the Cl_Bt regions. However RSG was not exclusive at any region, demonstrating concurrence and overlap at Cl_Bt regions. This significance and specificity of this data, at the multi-cellular level raises the question of a possible “bioelectric code” present and operative as part of the overall paleochronic reversion dynamic (Levine and Martyniuk, 2017).

RSG on the pericladial stalk and pedicel indicates that the putative morphogen(s), or signal(s) (Roberts and Friml, 2009) responsible for rhizogenous activity is not confined to specified floral organs, neither in presence nor in function but also can have effect in fields of expanded anatomic regions. Distinction between organ(s) RSG, zonal RSG and organs regions RSG become more apparent. RSG and variation in its site distribution are significantly associated with the calyx and bracts organs. That variation presents a vectorial cadastral localization in and through those organs. Results indicate definite regional homeostatic canalization and organ vectorial cadastral functions with eigenvector dynamics not only after paleochronic floral reversion occurred (Benya, 2001) but also associated with floral axial permutation (Benya and Windisch, 2007) whose specificity, degree and intensity affected succeeding and/or simultaneous RSG. The RSG function is robustly canalized. That canalization presents both linear and nonlinear (i.e. quadratic and/or cubic) aspects in its distribution (Green *et al.* 2017).

Significant variation in site activity at the bracts regions raises questions concerning their function(s). In non-reverted flowers, bracts may be a zone of transition from the photosynthetic, aerial region of non-sexual and indeterminate growth to the non-photosynthetic region of sexual reproduction and determinate growth. However a unique, perhaps transformational, referential and/or defining function seems to be manifest by the bracts and IBS on reverted flowers. Continuity of RSG at the pedicel both prior to and after floral whorl axial permutation suggests a distinction of this anatomic region from the whorls and bracts.

These results support a hypothesis that homeostatic canalization (Okamuro *et. al.* 1993) may be of more than one cadastral nature (Specchia et. al. 2010) raising the prospect of more than one generative phase or of various aspects to one unique generative phase. Biophysical functions affect both permutative and generative phases at the anatomic and morphologic regions studied here. Both appear to be distinct and mutually exclusive, and a certain hierarchy of priority is evident.

## Supplementary data

Figure 1: “map”: Linear sequence of the 74 RSG eigenvector loci (present = +; absent = 0) distributed over the 49 specimens in clusters of 1 to 3 specimens each by permutation distance according to PCL and/or IBS lengths (0.0 to 38.0 mm).

## ACKNOWLEDGEMENTS

Saint John’s College, Landivar, Belize City, Belize, C.A. (T. Thompson, professor) provided an introduction to material. G.R. Lovell (USDA-ARS Griffin, Gerogia, USA), W. Denny (USDA-ARS Beltsville, Maryland, USA), T.N. Khan, Dept. Agr. Western Australia and H.P.N. Gunasena, U. Peradeniya, Sri Lanka provided seed. A.C. Machin assisted with seed importation. Escola Agrícola Santo Afonso Rodriguez, (J. Moura Carvalho, E.M. Moreira, J. Bulfoni and I. Govoni) and Escola Técnica Soinho provided time and facilities to pursue these experiments. The “Universidade do Vale do Rio dos Sinos” (UNISINOS) and “Instituto de Pesquisa de Planarias” (IPP, Ana Leal-Zanchet, coordinator) furnished facilities for analysis of data. D. Nevins and P G. Windisch read and made important suggestions to improvement of the manuscript. A. DePaula, J. M. daSilva, E.O. Alves, J. deFreitas, C.G. deOliveira, D. Nevins, F. Gil, E.L. Confortin, A. Bruckschen, E. Loechner and G. & H. Galik helped with technical work and analysis. M.C. Moura Carvalho, C. Radz, M. Sander, S.J.V. Benya and T. H. Oliveira assisted with manuscript preparation.

## Compliance with ethical standards

**Conflict of interest:** The author declares that he has no conflict of interest.

This research did not receive any specific grant from funding agencies in the public, commercial or not-for-profit sectors.

**Novel material:** Seed, homozygous for the *srs* recessive allele (Benya and Windisch, 2007; Benya, 2016) is available upon request.

